# The Agroforestry Species Switchboard, a global resource to explore information for 107,269 plant species

**DOI:** 10.1101/2025.03.09.642182

**Authors:** Roeland Kindt, Ilyas Siddique, Ian Dawson, Innocent John, Fabio Pedercini, Jens-Peter Lillesø, Lars Graudal

## Abstract

The Agroforestry Species Switchboard is a comprehensive vascular plant database that guides users to information for a particular taxon from a global but fragmented set of resources. Via standardized species names, a user can rapidly infer which among the 59 contributing databases contain information for a species of interest, and understand how this information can be accessed. By providing taxonomic identifiers to World Flora Online, it is straightforward to check for changes in taxonomy. Among the 59 databases referenced, ten covered over 10,000 species, 20 between 1,000 and 10,000 species, and 22 between 100 and 1,000 species. The ten species-richest plant families across covered databases were the Fabaceae (9,537 species), Asteraceae (6,041), Rubiaceae (4,812), Poaceae (3,947), Myrtaceae (3,544), Euphorbiaceae (2,689), Malvaceae (2,478), Rosaceae (2,374), Lauraceae (2,334) and Lamiaceae (2,107). Information included in the Switchboard distinguishes 54,812 tree-like species, covering most known tree species globally. Among its applications, the Switchboard can assist species selection for ecological restoration projects to synergize biodiversity and human well-being objectives.

## Background & Summary

The earth’s life support systems depend on biodiversity to an extent that requires nature conservation far beyond protected areas (Van Der Plas et al. 2016; Dinerstein et al. 2020; Guo et al. 2023; Lima et al. 2023). Ecological restoration seeks to recover biodiversity and ecosystem integrity, while delivering ecosystem services and ensuring human well-being (Brancalion et al. 2017; Di Sacco et al. 2021; Kindt et al. 2023; Bartholomew et al. 2024; FAO, SCBD & SER 2024). Nature-based solutions implemented in human-modified landscapes are increasingly critical for social-ecological resilience (Lavorel et al. 2022; Dunlop et al. 2024). Yet taxon-specific knowledge that can inform restoration and diversification planning remains highly fragmented among dozens of ‘siloed’ databases, each with substantial gaps, contributing to the dominant use of a narrow set of intensely researched and promoted species in agriculture (Dawson et al. 2019), forestry (Pancel 2014) and sometimes even ecological restoration (Holl et al. 2022; de Almeida et al. 2024).

The Agroforestry Species Switchboard (hereinafter: the Switchboard) was specifically developed to facilitate access to detailed yet fragmented information for a broad range of tree species to support their wider use. The adoption of the term “Switchboard” is because of the role of the database as a central hub to guide users to a wide range of particular information sources. Its principal focus is on international databases that document tree species, supplemented with different types of database covering botanical, biochemical, biophysical, physiological, ecological, biogeographic, conservation, agronomic, silvicultural, nutritional, socioeconomic and cultural characteristics for plant taxa of all growth forms. Including 107,269 accepted plant names, the Switchboard in its current version spans over a quarter of accepted plant species globally and covers 59 scientific- and practitioner-oriented databases.

The Switchboard can be used by practitioners, researchers and policy makers to readily access descriptions of vascular plant species, including information on products and services provided, details on risks of invasiveness, and descriptions such as wood density and food nutritional compositions. Referencing of a species through the Switchboard to the included specialized databases such as on timber, forage and food uses can aid in prioritizing species for these functions, while information obtained through databases linked to the Switchboard related to species’ environmental ranges can further support species-site-use matching.

As linking through the Switchboard is done via a standardized plant name, users do not need to comb and check each individual reference database for potential synonyms or taxa with different spellings. An additional benefit, based on listing the original names of taxa used by databases, is that practitioners that are not familiar with recent taxonomic changes can recover information for taxa by their synonyms. Having standardized the great majority of taxa to the taxonomic backbone of World Flora Online (WFO; Borch et al. 2020), via the taxonomic identification field (‘taxonID’) users can rapidly retrieve data from the WFO website (https://www.worldfloraonline.org/), including information from taxonomic revisions, from particular flora, from the IUCN Red List of Threatened Species (https://www.iucnredlist.org/), and from distribution maps derived via the Plants of the World Online (https://powo.science.kew.org/).

Developers of new decision-support tools can improve the utility of their resources by linking to numerous other databases using the Switchboard as an intermediary. A recent published example of such an application is the GlobalUsefulNativeTrees database (GlobUNT; Kindt et al. 2023), which provides lists of native tree species for a selected country. Filtered species lists from GlobUNT link directly to entries in the Switchboard, which allows further narrowing down and the corroboration of species selections.

Given copyright statements and out of respect and in appreciation of the tremendous efforts made by those that compiled the information gathered in the constituent linked databases of the Switchboard, our intention is that users are directed to “fair usage”, recognition and appropriate citation of the contributing databases themselves.

## Methods

### Data collation

Here we document the collation of information for the Agroforestry Species Switchboard. Initial development of the Switchboard began in 2013 and it has now evolved to the present Version 4.0 which has reached a stage where its functionality merits wider adoption and promotion. The first criterion to link a database through the Switchboard is that it contains information about tree species, since diversifying tree use in restoration and broader tree planting initiatives is a primary concern for the authors and the institutes where they work. However, databases do not need to exclusively relate to trees and, where available, links to data on other vascular plant forms within the constituent referenced databases are included in the Switchboard. The second criterion to reference a database is that the information it contains has international, where possible global, coverage. However, databases focusing on larger nations that include different ‘botanical countries’ (level 3 areas in Brummitt’s 2001 *World Geographical Scheme for Recording Plant Distributions*) such as the USA (e.g., USDA-Plants [USDA 2024]) or Australia (e.g., EUCLID [Slee et al. 2020]) are included if these document sufficient numbers of species that are also grown more globally. Third, it should be possible to retrieve taxon-specific information easily from the database, in order to allow linkage via species names. Fourth, along with databases *per se*, a smaller number of resources linked to the Switchboard are books, scientific articles or spreadsheet documents where useful information is included that can easily be cross-linked within a database environment. Fifth, ideally the databases listed should be available at no cost to the user, although in certain cases exceptions were made. Supporting information in the *Switchboard contributing databases*.*xlsx* file (hereinafter: *metadata*) in the Zenodo archive indicates when reference database links were integrated into the Switchboard and how this was achieved, and provides further information on the properties of the individual linked databases. Future versions of the Switchboard will explore the integration of further databases.

Databases referenced within the Switchboard fall into four categories (Table 1, *metadata*).

1. ICRAF databases: these are databases that were (co-)developed by CIFOR-ICRAF. These include: the African Wood Density Database (abbreviation used: AWDD; Carsan et al. 2012), the Agroforestree Database (AFD; Orwa et al. 2009), the CIFOR-ICRAF Tree Genebank (ICRAF-gene; CIFOR-ICRAF 2024), GlobalUsefulNativeTrees (GlobUNT; Kindt et al. 2023), the Priority Food Tree and Crop Food Composition Database (ICRAF-nutri; Stadlmayr et al. 2019), RELMA-ICRAF Useful Trees (RELMA-ICRAF; Kindt & Innocent 2016), TreeGOER: Tree Globally Observed Environmental Ranges (TreeGOER; Kindt 2023a), Useful Tree Species for Africa (UTSA; Kindt 2023b) and the Vegetation map for Africa species distribution maps (V4A; Kindt et al., 2015).
2. Invasive species databases: these are databases that document potentially invasive species (Pyšek et al. 2020). Identifying invasive species – in order not to introduce them and to remove them if present – can be a key process in ecological restoration that is explicitly mentioned in ecosystem restoration principles (Bartholomew et al. 2024; FAO, SCBD & SER 2024). For this reason and also after a suggestion from the Deputy Executive Secretary at the Convention of Biological Diversity Secretariat on how to feature these species, these databases are listed in a separate category. These databases include: CABI Compendium Invasive Species (CABI-ISC; CABI 2022a), the Country Compendium of the Global Registry of Introduced and Invasive Species (GRIIS; Pagad et al. 2018) and the Global Invasive Species Database (GISD; ISSG 2024).
3. Spreadsheet databases: these are databases that typically can be downloaded as a single file, where the user can find information for the taxon by searching for its name within the document. Some databases can be searched online, but we did not obtain taxon-specific links. Some databases could be searched online previously. These include: A global database of plant services for humankind (GPS; Molina-Venegas et al. 2021), Árboles de Centroamérica: un Manual para Extensionistas (Spanish) (AdC; Barrace et al. 2003), BIOMASS: Estimating Aboveground Biomass and Its Uncertainty in Tropical Forests (BIOMASS; Réjou-Méchain et al. 2017), Dr. Duke’s Phytochemical and Ethnobotanical Databases (Duke-Ethno and Duke-Phyto; USDA 2016), eHALOPH: a database of halophytes and other salt-tolerant plants (eHALOPH; The University of Sussex 2024); Especies para la restauración en Mesoamérica (Spanish) (Restoracion; Sanchún et al. 2016), the Food Plants International database (FoodPlants; Food Plants International 2024), GlobaAllomeTree: Assessing volume, biomass and carbon stocks of trees and forests (GlobAllome; Henry et al. 2013), the Global Species Matrix (GSM; Toensmeier 2016), the Tropical Forestry Handbook: Species Files in Tropical Forestry (TFH; Pancel 2015), the Tallo database (Tallo; Jucker et al. 2022), The International Timber Trade: a working list of commercial timber tree species (ComTimber; Newton et al. 2014), the World Checklist of Useful Plant Species (WCUPS; Diazgranados et al. 2020) and the World list of plants with extrafloral nectaries (Extrafloral; Keeler et al. 2024).
4. Other databases: these are databases not listed in the previous categories and for which taxon-specific links were included. These include: the African Orphan Crops Consortium priority crops (AOCC; AOCC 2017), Australian Tropical Rainforest Plants (LUCID-rain; Zich et al. 2020), CABI Compendium Forestry (CABI-Forest; CABI 2022b); the CABI Compendium (CABI; CABI 2024), ECHOcommunity Plant Search (ECHO; ECHOcommunity 2022), the ECOCROP Database of Crop Constraints and Characteristics (ECOCROP; FAO 2013), EUCLID Eucalypts of Australia (LUCID-euc; Slee et al. 2020), European Forest Genetic Resources Programme Species (EUFORGEN; EUFORGEN 2024), FamineFoods (FamineFoods; Freedman 2024), Feedipedia (Feedipedia; INRAE, CIRAD, AFZ and FAO 2024), the INBAR Bamboo and Rattan Species Selection tool (INBAR; INBAR 2022), ITTO Lesser Used Species (TropTimber; ITTO 2024), Mansfeld’s World Database of Agriculture and Horticultural Crops (Mansfeld; Mansfeld 1959), the New World Fruits Database (NWFD; Bioversity International 2017), NewCROP (NewCROP; Purdue University 2022), Optimizing Pesticidal Plants (OPTIONs; Stevenson 2014), Oxford Plants 400 (Oxford400; UoO 2022), Plant Resources of South-East Asia (PROSEA; Verheij et al. 2022), Plant Resources of Tropical Africa (PROTA; Grubben et al., 2022), the PLANTS Database (USDA-Plants; USDA 2024), Plants For A Future (PFAF; PFAF 2024), Portal de Plantas Medicinais, Aromáticas e Condimentares (Portuguese) (PPMAC; PPMAC 2024), Seed Leaflets (Leaflets; UoC 2024), Species Profiles for Pacific Island Agroforestry (Pacific; Elevitch 2006), The Gymnosperm Database (Gymnosperm, Earle 2024), The Wood Database (TheWood; Meier 2024), Tropical Forages (SoFT; Cook et al. 2020), tropiTree (TropiTree; Russell et al. 2014), Useful Temperate Plants (UTempP; Fern 2024a), Useful Tropical Plants (UTropP; Fern 2024), WATTLE Acacias of Australia (LUCID-wat; Maslin 2018) and World Economic Plants in GRIN-Global (USDA-GRIN; Wiersema & León 1999).

**Table 1.**
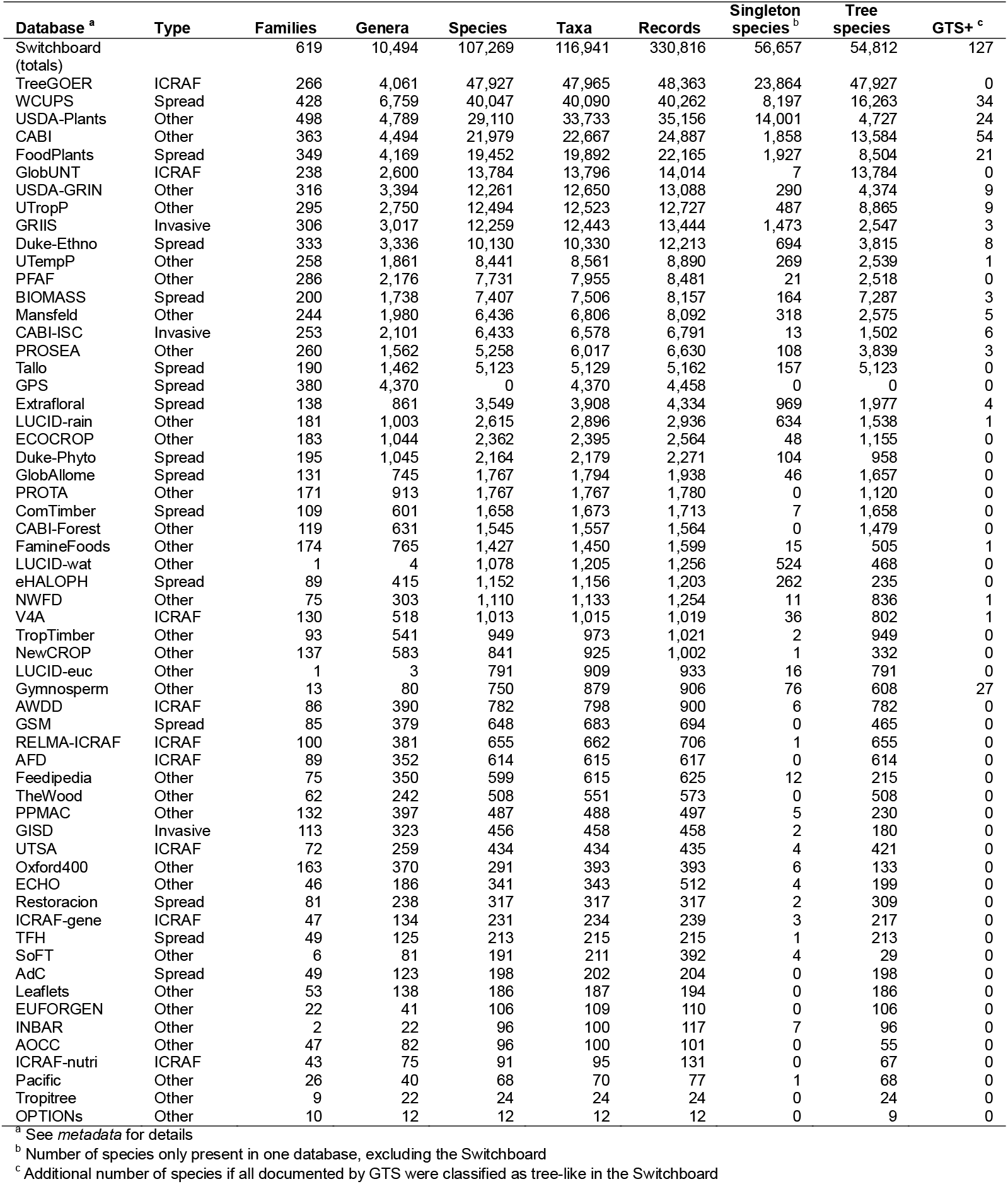
The 59 contributing databases referenced in the Switchboard and the taxonomic diversity represented. Databases are sorted by overall taxon richness.

### Taxonomic standardization

Taxonomic standardization among referenced databases and the Switchboard was achieved in the *R* statistical environment (version 4.2.1; R Core Team 2022; figures included here were generated in the same environment) with functions WorldFlora::WFO.match.fuzzyjoin() and WorldFlora::WFO.one() from the WorldFlora package (version 1.14-5; Kindt 2020). For databases where information was downloaded on the naming authority (see *metadata*), this information was also utilized during matching via the ‘Authorship’ argument, similar as in workflows documented by Kindt (2023c). Taxonomic names were matched first against the taxonomic backbone of WFO (<u>December 2023 Release</u>). Direct matches retained in subsequent standardization steps were documented as ‘direct’ in the ‘Match.Type’ field; the Switchboard database (archived as: *Switchboard_4*.*txt*) contains 318,268 entries of this match category. Suggested fuzzy matches accepted after visual inspection were documented as ‘manual’ in the ‘Match.Type’ field; the Switchboard contains 12,408 retained entries of this match category. Next, still unmatched names were compared with the taxonomic backbone of the World Checklist of Vascular Plants (WCVP; version 11; Govaerts et al. 2021). Where matches here were identified, these were assigned to ‘direct’ or ‘manual’ categories, referencing ‘WCVP’ in the ‘Backbone’ field; the Switchboard includes 202 and 97 such entries, respectively. A final step involved the matching of current names suggested for synonyms via WCVP with the WFO backbone. Where matches here were indicated, these were documented as ‘direct via WCVP’ or ‘manual via WCVP’ entries (93 and 47 records, respectively).

The taxonomic standardization described above was achieved by processing the contributing databases individually, except for two groups of databases that were processed collectively. The first among these groups (labelled as ‘STATIC’ databases, see *metadata*) included databases where contents had not changed since the third version of the Switchboard, or where we failed in obtaining a more current taxon-based data set. The second group of databases (‘ZENODO’ databases, see *metadata*) were databases accessed from Zenodo archives. Both STATIC and ZENODO databases had been standardized earlier with similar standardization protocols as described above, but with an older version of the WFO taxonomic backbone. When processing the STATIC and ZENODO databases, we first matched species names via the WFO ‘taxonID’ field from the match with the older version of WFO with the ‘taxonID’ from the more recent version used in standardization workflows for Switchboard 4.0, provided that the older matches had been classified as ‘direct’. Still unmatched taxa were then processed further with the same WorldFlora workflows applied to individual databases.

Taxa not matched by the above approaches were removed from the Switchboard if identified as taxonomic names for algae (the main resource for screening was https://www.algaebase.org/), fungi (comparing with https://www.indexfungorum.org/names/names.asp), bacteria or animals (confirmed via online searches). The *metadata* indicates for which of the 59 registered databases such organisms were encountered and provides examples. Also removed were cultivar entries for plants such as *Actinidia arguta* ‘AADA’, and hybrids, where databases listed them in a format such as *Megathyrsus maximus* x *M. infestus*.

Several taxa at infraspecific levels (var., subsp. and f.) could not be matched with these procedures. These infraspecific taxa were matched next with a previous species-level match. Where this was not possible as there was no previous entry at species-level, we conducted the standardization protocols at species level.

When the above standardization steps were completed, we noticed situations where taxa of the same taxon name had been matched with different records in the taxonomic backbone database. Typically, this was a consequence of having authorities for taxa available in some databases but not in others. To deal with these situations – unless there was information on the authority from the database that suggested otherwise – in the case of trees, we decided to select the same match that was made with the full list of 57,681 species names and their naming authorities from GlobalTreeSearch version 1.8 (GTS; Beech et al. 2017; we did not include GTS information in Switchboard 4.0 for copyright reasons, hence selecting the same matches also facilitates comparison of the two databases). In a next series of updates based on the same procedure, we, where possible, resolved matches for species not covered by GTS by adopting the match made with the 13,088 World Economic Plants in GRIN-Global database (Wiersema & León 1999), which provides naming authorities. For practical reasons, standardizations made via GTS and USDA-GRIN databases were done first for different batches of databases (see *metadata*) and then repeated after combining information from all databases.

After concluding the workflows described above, we remained with 1,169 records, potentially listing an additional 1,109 taxa for the Switchboard, that could not be matched and were therefore not included in the *Switchboard_4*.*txt*. The list of unmatched taxa is available as a separate data file *Switchboard_unmatched*.*txt*.

The final checks to identify and correct situations where taxa had been matched differently, when there was information that in fact these were the same taxa, are described in the section on Technical validation below.

### Taxonomic, geographic and phylogenetic coverage

The Switchboard contains 619 families, 10,494 genera and 107,269 species of vascular plants (Table 1). Two databases covered the highest numbers of standardized species, TreeGOER (47,927) and WCUPS (40,047). These were among ten databases that covered over 10,000 species. 20 databases covered between 1,000 and 10,000 species, and 22 between 100 and 1,000 species. Seven databases covered less than 100 species, but these included the GPS that provides data at the genus level for 4,370 genera.

We inferred the continental native distribution of species from information in the WCVP. The WCVP identification field (‘plant_name_id’) to obtain distribution on geographical distributions was inferred by linking the WFO and WCVP backbone databases via their ‘IPNI ID’ fields. This was achieved for 96,957 species listed in the Switchboard. Where such link could not be established by matching IPNI identification fields, we matched species via WorldFlora and the WCVP taxonomic backbone, accepting 8,168 matches. The matched WCVP identification (‘WCVP_ID’) is included in the species file of the Switchboard.

For four continents (Africa [AFR], tropical Asia [OAS], Southern America [SAM] and Australasia [AUS]), the two most species-rich databases (TreeGOER and WCUPS) had the highest species richness (Supplementary Table 1). For temperate Asia [EAS], FoodPlants ranked second highest, for northern America [NAM] USDA-Plants had the highest richness. USDA-Plants ranked second highest both for the Pacific [PAC] and Europe [EUR], whereas GRIIS had the highest richness for Europe. 54 databases contained species native to 7 or all continents. EUR was the continent with fewest database coverage, although there still 48 databases containing some of its native species.

The ten species-richest plant families were the Fabaceae (9,537 species), Asteraceae (6,041), Rubiaceae (4,812), Poaceae (3,947), Myrtaceae (3,544), Euphorbiaceae (2,689), Malvaceae (2,478), Rosaceae (2,374), Lauraceae (2,334) and Lamiaceae (2,107) (Supplementary Table 2), collectively including 37.2% of species of the Switchboard. Excluding the Myrtaceae from the six species-richest families, these were the same families identified as the most diverse in the WCUPS (Diazgranados et al. 2020). Across their native continents, the five most frequent families included those with the highest and second-highest richness. The Fabaceae were the richest families for species native to AFR, OAS, SAM and AUS, the Asteraceae for SAM and EUR, the Rubiaceae for PAC and the Poaceae for EAS.

The cladogram (Figure 1) summarizes the phylogenetic diversity of the Switchboard. The phylogenetic tree was constructed via the V.PhyloMaker2 R package (version 0.1.0; Jin and Qian 2022) with the GBOTB.extended.TPL tree and “S3” scenarios. To represent families, we selected the most frequent species for each family in the Switchboard and matched these with the same species, same genus or another species of the family of the GBOTB tips (Supplementary Table 2 documents the matched species and taxonomic level of matching). 177 families were excluded from the analysis as no matching GBOTB tip was found. The cladogram was constructed via the ggtree (version 3.6.2; Yu 2023) and ggtreeExtra (version 1.8.1; Xu et al. 2021) packages with ggplot2 (version 3.5.1; Wickham 2016;). The cladogram depicts the wide taxonomic and geographic coverage of the Switchboard, such as 106 families being native to all continents and 67 families being native to a single continent.

**Fig. 1:**
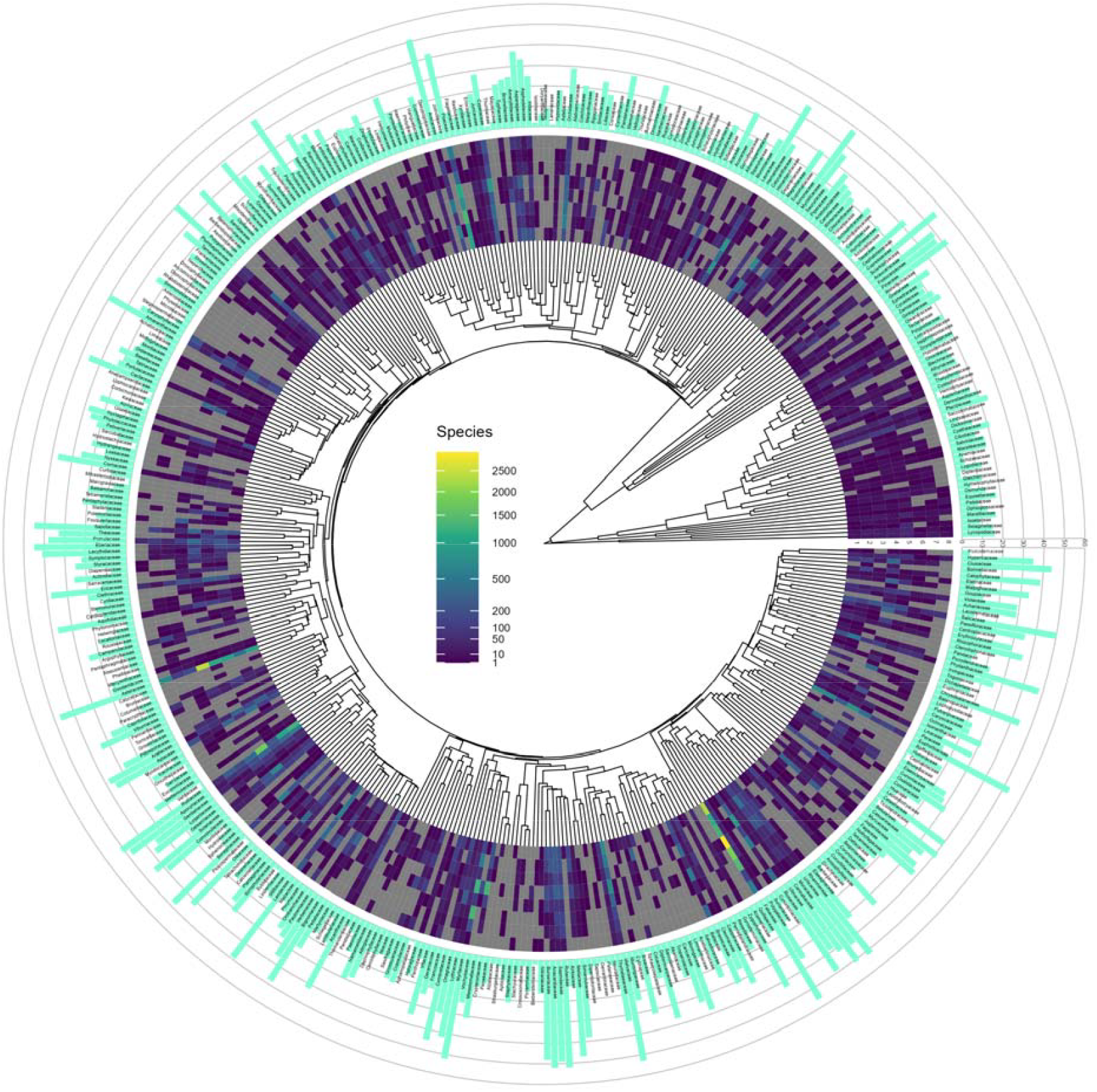
Cladogram of 442 plant families documented in the Switchboard. The circular heatmap shows the number of species native to Africa (1), tropical Asia (2), temperate Asia (3), southern America (4), northern America (5), Australasia (6), Pacific (7) and Europe (8) based on the WCVP. The bar graphs show the number of databases with information about a family.

### Flagging of tree species

Not including the global species list from GTS in Switchboard version 4.0 due to copyright restrictions presented the opportunity for an alternative system of identifying species as tree or woody perennial species. We opted for a classification system that also potentially included taxa that had been deliberately excluded from GTS of cycads, tree ferns, tree-like Poaceae, Bromeliaceae, Musaceae and hybrid species (Beech et al. 2017). For example, *Leucaena leucocephala* no longer features in GTS as it was hypothesized that the species arose as a direct result of cultivation of one or both of its diploid parents (personal communication with Emily Beech, head of Conservation Prioritisation, BGCI, 6^th^ September 2024). GTS adopted the tree definition agreed by IUCN’s Global Tree Specialist Group: a woody plant with usually a single stem growing to a height of at least two metres, or if multi-stemmed, then at least one vertical stem five centimetres in diameter at breast height.

The species file for TreeGOER provides the field of ‘Tree’ that flags and defines 54,812 tree species *sensu* the Switchboard. The file includes in a separate field (‘Tree-like justification’) containing the justification for flagging a species as a tree species via a consecutive process whereby species not previously flagged as trees were checked.

1. Batch 1 flagged species included in TreeGOER (47,927 species), TreeGOER+ (2,031; Kindt 2024a), GlobUNT (1,004) or the species list from an article providing a synthesis of global tree biodiversity data (1,229, the same standardization pipeline was applied to the appendix of this article; Keppel et al. 2021).
2. Batch 2 flagged species not identified in batch 1 with the lifeform described in the WCVP of liana (1,576 species), tree (651), herbaceous tree (92), bamboo (25) or herbaceous bamboo (4).
3. Batch 3 flagged 179 species present in Tallo (a database described as containing records of individual trees; Jucker et al. 2022) and that were not flagged previously.
4. Batch 4 flagged species present in databases expected only to include tree species given the scope of these databases: RELMA-ICRAF (43 species), AWDD (4), INBAR (4), ComTimber (2), AFD (1), EUFORGEN (1), Pacific (1), TheWood (1) and TropTimber (1).
5. Batch 5 flagged species that had not been flagged earlier, where there was information in a database included in the Switchboard that aided their flagging as trees: UTropP (13 species), PROSEA (10), LUCID-rain (4), USDA-Plants (2) and UTempP (1).
6. Batch 6 flagged five species that had not been flagged earlier, where there was information available in WFO of a tree-like description.

This flagging process in effect represents one application domain for the Switchboard, where information on presence of a species in a database allows specific analyses. Another direct application that is not comprehensively shown here is to rank and tentatively prioritize (see Kindt et al. 2021) species by the number of databases where they occur. The 25 most frequent tree species in the Switchboard are *Tamarindus indica* (40 databases), *Samanea saman* (39), *Vachellia nilotica* (39), *Cocos nucifera* (38), *Mangifera indica* (38), *Albizia lebbeck* (37), *Ceiba pentandra* (37), *Anacardium occidentale* (36), *Neltuma juliflora* (36), *Olea europaea* (36), *Psidium guajava* (36), *Eucalyptus camaldulensis* (35), *Gliricidia sepium* (35), *Persea americana* (35), *Azadirachta indica* (34), *Faidherbia albida* (34), *Melia azedarach* (34), *Moringa oleifera* (34), *Senna siamea* (34), *Acacia auriculiformis* (33), *Cedrela odorata* (33), *Enterolobium cyclocarpum* (33), *Jatropha curcas* (33), *Leucaena leucocephala* (33) and *Morus alba* (33).

### Taxonomic, phylogenetic and geographic coverage of tree species

The 10 most species-rich families for tree species were the Fabaceae (5,061 species), Rubiaceae (3,954), Myrtaceae (3,353), Lauraceae (2,304), Euphorbiaceae (1,823), Malvaceae (1,692), Melastomataceae (1,552), Annonaceae (1,481), Arecaceae (1,332) and Sapotaceae (1,109) (Supplementary Table 2). These were the same fop 10 families identified by Beech et al. (2017), whereas we also observed the same ranking by species richness. Across native continents, the top-4 families globally included the richest and second-richest families for AFR, OAS, SAM, AUS and PAC. However, the Rosaceae that globally ranked 12^th^ with 1,082 tree species was the richest in EAS (417 species) and EUR (176). In NAM, the Fagaceae were the second-richest with 280 species, a family that globally ranked 16^th^ with 860 species. Poaceae and Musaceae species excluded from GTS ranked 23^rd^ (617 species) and 101^st^ (65 species), respectively. Bromeliaceae species that were excluded from GTS were also not included among trees sensu the Switchboard, whereas among all species this family ranked 110^th^ (177 species, including *Ananas comosus* as the most frequently encountered across databases). The Switchboard tree species include 24 Zamiaceae (among all species in the switchboard, there were 138 species), 13 Cycadaceae (overall, 32) and 20 Cyatheaceae (overall, 100) species, where cycads and tree ferns had been excluded from GTS (Beech et al. 2017).

The cladogram for tree species represents all 285 families (Fig 2.)

**Fig. 2:**
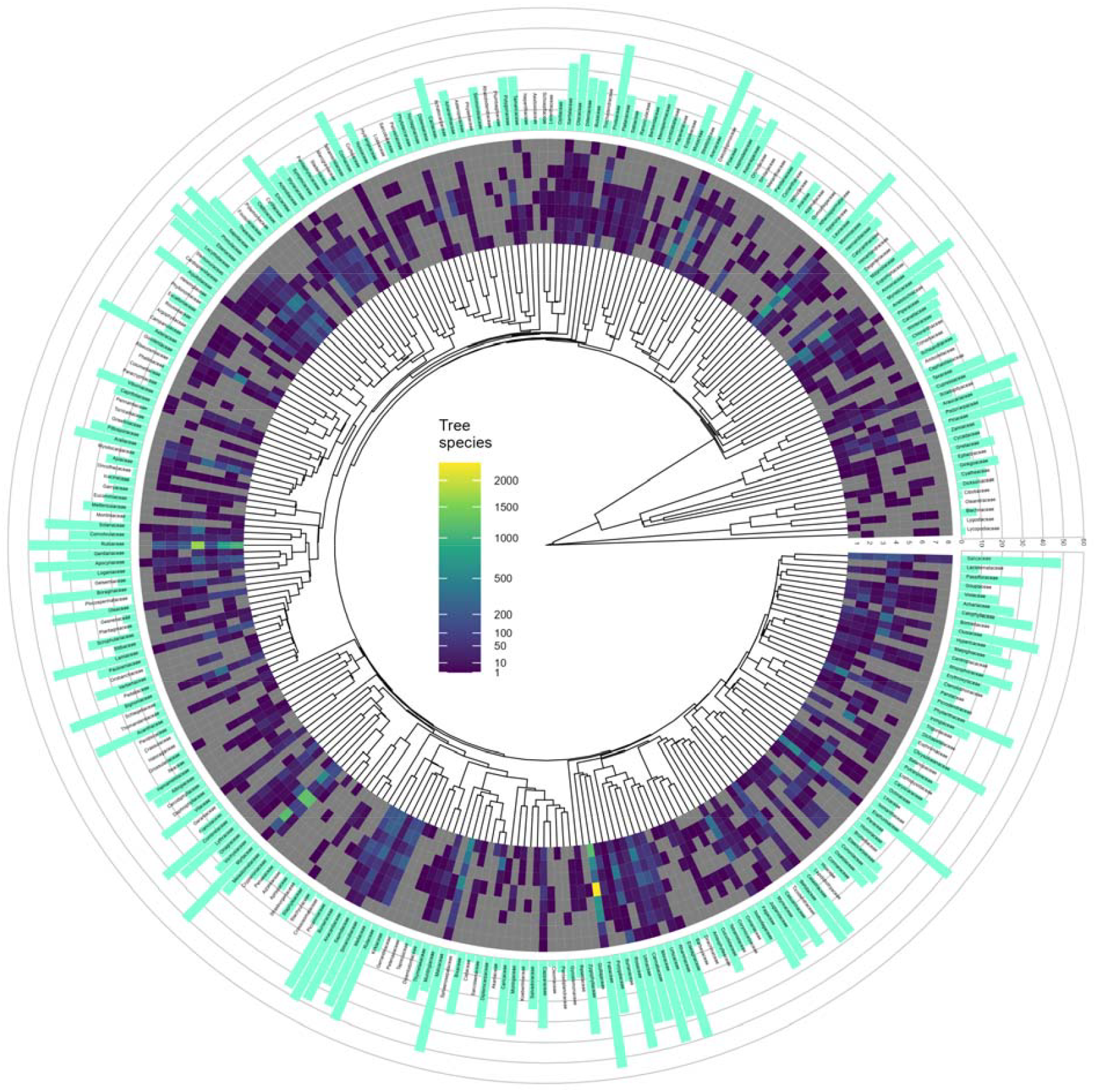
Cladogram of all 285 plant families containing tree species documented in the Switchboard. The circular heatmap shows the number of tree species native to Africa (1), tropical Asia (2), temperate Asia (3), southern America (4), northern America (5), Australasia (6), Pacific (7) and Europe (8) based on the World Checklist of Vascular Plants. The bar graph shows the number of databases with information about a family.

The dendrogram showing the relationship in tree species composition between databases (Figure 3) was constructed by first creating a distance matrix showing the pairwise Simpson dissimilarity measure (β_sim_; this index efficiently discriminates species turnover from nestedness; Baselga 2009), conducting hierarchical clustering with ‘average’ distance and reordering the tree by species richness of the databases. These calculations were achieved via the BiodiversityR (version 2.16-1; Kindt & Coe 2005), stats (version 4.2.1; R Core Team 2022) and vegan (version 2.6-4; Oksanen et al. 2024) packages. The dendrogram was constructed via the ggtree (Yu 2023, version 3.6.2) and ggtreeExtra (Xu et al. 2021; version 1.8.1) packages with ggplot2 (Wickham 2016, version 3.5.1), with similar scripts as the cladograms of figures 1 –2.

**Fig. 3:**
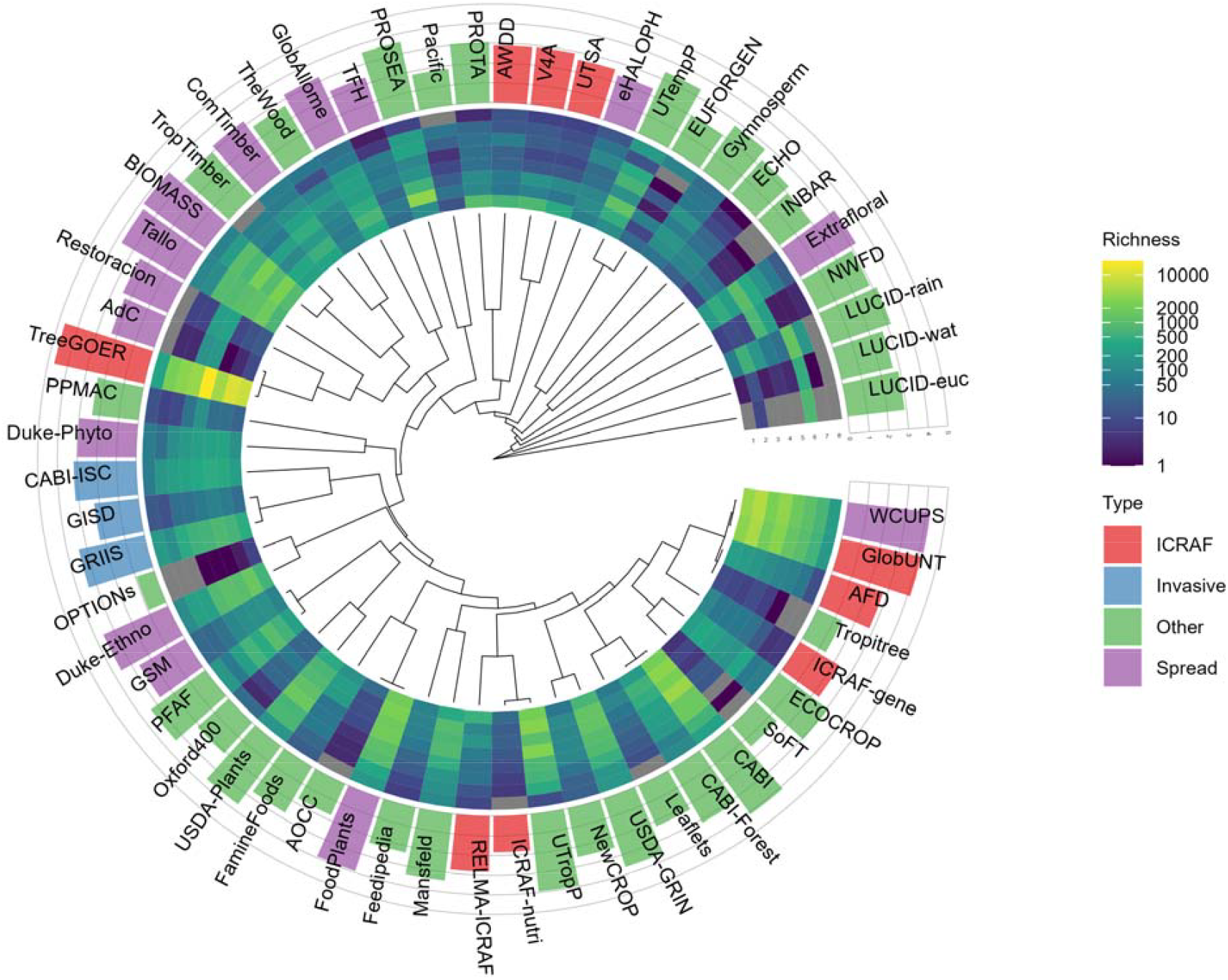
Clustering tree showing similarity in tree species composition (β_sim_) among databases documented in the Switchboard. The circular heatmap shows the number of tree species native to Africa (1), tropical Asia (2), temperate Asia (3), southern America (4), northern America (5), Australasia (6), Pacific (7) and Europe (8) based on the WCVP. The bar graph shows the number of tree species on a log10 scale.

Several patterns can be observed in species turnover. The three invasive species databases form a separate cluster. Databases focusing on specific continents also tend to cluster together such as the Africa-focused PROTA, AWDD, V4A and UTSA, or the European-focused EUFORGEN and temperate-focused UTempP. As expected by the way GlobUNT was created partially from information available in WCUPS, these two database cluster together. A similar reason explains the clustering of the GSM and PFAF databases. Databases focused on human food also cluster together, shown by the clustering of the AOCC, FoodPlants and FamineFoods databases. Databases containing information on timber species or wood density cluster first, shown by the pattern for the BIOMASS, TropTimber, TheWood, GlobAllome and TFH. TreeGOER clusters first with AdC, an indication that most species of the latter are included in the former, but in later clustering steps then clusters together with the timber and wood density cluster. Databases for specific taxa from specific regions only join the other clusters at final steps, as shown for the LUCID databases.

## Data records

The Switchboard can be downloaded as text files from a Zenodo archive under a CC-BY 4.0 License (https://doi.org/10.5281/zenodo.14991887; see Table 2). These text files where fields were separated by the pipe ‘|’ character can be directly imported in statistical software such as R, or pasted into spreadsheets. Also included in the archive is a spreadsheet in MS Excel format that provides metadata for the different databases covered by the Switchboard.

**Table 2.** Content of text file downloads available from the Agroforestry Species Switchboard archive.

| File | Field name | Notes |
| --- | --- | --- |
| Switchboard_4 | Database | Database code, see Databases.xlsx spreadsheet |
|  | Taxon_Database | Taxon name in the database |
|  | Taxon_URL | Taxon-specific part of the hyperlink |
|  | Match.Type | Method of matching names (see manuscript) |
|  | Backbone | Taxonomic backbone: WFO = World Flora Online; WCVP = World Checklist of Vascular Plants |
|  | SID | Taxonomic ID from the taxonomic backbone |
|  | ScientificFinal | Standardized taxonomic name |
|  | ScientificNameAuthorship | As in the taxonomic backbone |
|  | TaxonomicStatus | As in the taxonomic backbone |
|  | TaxonRank | As in the taxonomic backbone |
|  | Genus | As in the taxonomic backbone |
|  | Family | As in the taxonomic backbone |
|  | Species | Derived from ScientificFinal, also omitting notations ('x') for hybrid species |
| Switchboard_species | Species | Unique Species from Switchboard_4 file |
|  | Databases | Number of databases |
|  | Tree | Identification as tree |
|  | Treeish_batch | Batch when species was identified as tree |
|  | Treeish_justification | Justification for identification as tree |
|  | SID | As in the Switchboard_4 file |
|  | WCVP_ID | Taxonomic ID in the World Checklist of Vascular Plants |
|  | WCVP_ID_match | Method of obtaining the WCVP_ID (see manuscript) |
|  | TRY_Species<sup>a</sup> | Number of records in the TRY harmonized species data set with the same SID |
|  | WFO_Accepted | Number of records in the WFO taxonomic backbone with the same species name and where the taxonomicStatus was 'Accepted' |
|  | WFO_Unchecked | Number of records in the WFO taxonomic backbone with the same species name and where the taxonomicStatus was 'Unchecked' |
|  | WFO_Synonym | Number of records in the WFO taxonomic backbone with the same species name and where the taxonomicStatus was 'Synonym' |
|  | WFO_none | Indicator that the same species name was not listed in the WFO taxonomic backbone. This indicator also flags 'Species' with a hybrid ('x') notation in the taxonomic backbone. |
| Switchboard_unmatched | Database | See Switchboard_4 |
|  | Taxon_Database | See Switchboard_4 |
|  | Taxon_URL | See Switchboard_4 |

<sup>a</sup> A list of plant names in the TRY Plant Trait Database (Schellenberger et al. 2023) was standardized to the same taxonomic backbone data via WorldFlora (Kindt 2024b). This dataset can be linked with the Switchboard via the 'SID' field.

## Technical validation

Besides the taxonomic standardization pipelines described above, in a series of manual checks after information for all databases was combined, we resolved various cases of taxa that were matched with different taxonomic records, but that could be matched with the same record:

- Taxa with different matching WFO ‘taxonID’, but with the same (including spelling variants) name, authority and IPNI in the WFO taxonomic backbone were cross-checked and manually resolved to a single record where possible. Examples included *Balanites aegyptiaca* (L.) Delile [accepted name; <u>wfo-0000313273</u>] and *Balanites aegyptiacus* (L.) Delile [accepted name; <u>wfo-0001146307</u>], *Acacia macdonnellensis* Maconochie [accepted name; <u>wfo-0000202850</u>] and *Acacia macdonnelliensis* Maconochie [accepted name; <u>wfo-0000202851</u>], and *Acalypha alopecuroidea* Jacq. [accepted name; <u>wfo-1000054518</u>] and *Acalypha alopecuroides* Jacq. [accepted name; <u>wfo-0000275337</u>]. We used the same protocols as described for the standardization procedures, giving preference to the names matched from GTS and USDA-GRIN. Where this was not possible, randomly one option was selected. Note, however, that the original names in the different databases are shown also in the Switchboard.
- Taxa of the same (including spelling variants) names that were first matched with different records that further included more than one taxonomic status between accepted, synonym and unchecked were manually matched with a single record. Examples included taxa first matched with *Acacia albida* Rojas Acosta [unchecked; <u>wfo-1200021925</u>] or *Acacia albida* Delile [synonym; <u>wfo-0000188059</u>], *Austroeupatorium inulifolium* (Kunth) R.M.King & H.Rob [unchecked; <u>wfo-1000056230</u>] or *Austroeupatorium inulaefolium* (Kunth) R.M.King & H.Rob. [accepted; <u>wfo-0000050339</u>], and *Sarcocephalus pobeguinii* Pobég. [unchecked; <u>wfo-0000303401</u>] or *Sarcocephalus pobeguinii* Hua ex Pobég [synonym; <u>wfo-0001225760</u>]. We resolved these cases by checking for the accepted name from the online (2024) WFO and POWO. For example, in the Switchboard we have treated *Acacia albida* as synonym of *Faidherbia albida*.
- Taxa with the same accepted name, but with different naming authorities according to different records were manually resolved by selecting one of the records, based on information available on authorities in the GTS or USDA-GRIN databases. This eliminated most cases, except for the following taxa with different records and authorities: *Calophyllum polyanthum* (<u>wfo-0000581286</u> with authority of Wall. ex Planch. & Triana; <u>wfo-0001296210</u> with Wall. ex Choisy), *Hippeastrum puniceum* (<u>wfo-0001046684</u> with (Lam.) Kuntze; <u>wfo-0000661183</u> with (Lam.) Voss), *Pluchea sagittalis* (<u>wfo-0000016241</u> with Less; <u>wfo-0000017448</u> with (Lam.) Cabrera, *Prunus emarginata* (<u>wfo-0001013841</u> with (Douglas ex Hook.) Walp.; <u>wfo-0001006712</u> with (Douglas) Eaton) and *Sorbus torminalis* (<u>wfo-0000998680</u> with Garsault; <u>wfo-000101407</u> with (L.) Crantz). For these five taxa, we included different taxonomic IDs in the main Switchboard file, but selected a single taxonomic entry (the first one shown in the previous sentence) in the species file. This process resulted in having a one-to-one match of species names with the taxonomic IDs in the species file.

The *Switchboard_species*.*txt* file contains fields that show the number of potential candidate direct matches between the resolved species name and the records of the World Flora Online backbone. Where multiple candidates exist, as for 7,865 (7.3%) of species, the user may wish to reconfirm the species-backbone matches, especially if this is possible with information on authorities. Similarly, where matching was done manually, as explicitly document by the Switchboard, users may wish to cross-check the name match.

## Usage notes

The main objective of the Switchboard is to show the user where information is available for a particular species, and how this information can be retrieved via the name used in the database. As taxonomy is a dynamic field, users interested in a particular taxon are encouraged to check for current names via WFO, which can be done via the ‘taxonID’ as shown in the previous section. Users could especially cross-check species that are listed as unchecked in the Switchboard ‘taxonomicStatus’ field, as those could have been resolved as current names or synonyms in newer versions of WFO or WCVP.

Information from the Switchboard can be used internally in decision support tools such as the GlobUNT database (Kindt et al. 2023). The *metadata* contains examples of how taxon-specific hyperlinks can be constructed to different databases. To act as a central hub for taxon-specific information, users can create an URL-friendly link to the Switchboard application by a naming convention such as https://apps.worldagroforestry.org/products/switchboard/index.php/name_like/Prunus%20africana, where the hyperlink connects to the online version of the Switchboard available via the CIFOR-ICRAF website.

## Acknowledgment

We are indebted to the tremendous efforts of experts that created the different databases covered by the Switchboard, as well as those that created taxonomic backbone data sets. We greatly appreciate the assistance provided by Christopher J. Earle in sharing a sitemap of the Gymnosperm database. CIFOR-ICRAF’s collaborative work with partners on trees, including on decision-support tools, receives significant financial support. Support for decision-support tools to enhance tree planting comes from Norway’s International Climate and Forest Initiative through its funding of the Provision of Adequate Tree Seed Portfolios project (PATSPO); from Germany’s International Climate Initiative through its funding for the Right Tree in the Right Place project (RTRP-Seed); from Darwin Plus (The Overseas Territories Environment and Climate Fund) through its funding of The Global Biodiversity Standard (Grant No. DAREX001); from the Bezos Earth Fund that invests in tree seed and seedling system development in Kenya and the Lake Kivu and Rusizi River Basin; and from the Green Climate Fund (GCF) through the IUCN-led Transforming the Eastern Province of Rwanda through Adaptation project (TREPA) and through the Climate Appropriate Portfolios of Tree Diversity in Burkina Faso (R-CAPTD). In addition, CIFOR-ICRAF gratefully acknowledges the support of the EU and broader CGIAR funding partners. I.S. thanks the Coordenação de Aperfeiçoamento de Pessoal de Nível Superior -Brasil (CAPES) -Finance Code 001 through PrInt-CAPES/UFSC of the Postgraduate Program in Plant Genetic Resources (RGV-UFSC).

